# Brassinosteroid-auxin crosstalk shapes rice flag leaf angle to boost dense planting yield

**DOI:** 10.64898/2026.07.30.741145

**Authors:** Suhang Yu, Qiaoyi Li, Yali Xiong, Dilixiadanmu Tashenmaimaiti, Zhuo Qu, Yang Yang, Jinge Tian, Guoqiang Huang, Xiuzhen Kong

**Author notes:** Co-corresponding author: Xiuzhen Kong and Guoqiang Huang. The author(s) responsible for distribution of materials integral to the findings presented in this article in accordance with the policy described in the Instructions for Authors (www.plantcell.org) is: Xiuzhen Kong and Guoqiang Huang.

## Abstract

Flag leaf angle (FLA) critically determines rice yield potential under dense planting conditions. As two pivotal phytohormones determine rice FLA, the antagonistic interaction between brassinosteroid (BR) and auxin remains largely uncharacterized. We here demonstrate that BR signaling reduces auxin biosynthesis to control FLA via regulating the biosynthesis of secondary cell wall (SCW). Genetic evidence demonstrates *OsYUC8* mutants disrupt SCW formation, leading to increased FLA. At the molecular level, the BR-related transcription factor OsBZR1 directly binds to and represses *OsYUC8* promoter activity, thereby fine-tuning auxin-mediated SCW biosynthesis. Field evaluations reveal that *osbzr1* mutants display optimized flag leaf architecture and improved yield performance under high-density cultivation. Our study not only delineates the antagonistic BR-auxin interaction governing FLA but also establishes a genetic strategy for manipulating crop architecture to maximize yield in dense planting conditions.

**One-sentence summary:** OsBZR1-mediated repression of *OsYUC8*-driven auxin biosynthesis modulates secondary cell wall formation to optimize flag leaf architecture, providing a genetic strategy for maximizing yield under high-density cultivation.

## Introduction

Flag leaf of rice, which is the topmost leaf closest to the panicle, is a crucial energy source for panicle development, making the optimization of its erectness is particularly important (Sakamoto et al., 2006). Upright flag leaf can receive light from both sides, improve light capture efficiency, reduce the obstruction of lower light, increase light transmission and enhance photosynthesis of lower leaves, which are more tailored for dense planting (Zhang et al., 2021; Cao et al., 2022; Xing et al., 2022). Lamina joint is the unique structure connecting leaf blade and leaf sheath, determining leaf angle by controlling the degree to which the leaf blade bends away from the leaf sheath (Liu et al., 2024). Auxin and BR, two crucial phytohormones, act antagonistically in regulating rice FLA, with auxin promoting leaf erectness while BR enhances leaf inclination.

Auxin negatively regulates rice leaf angle through modulation of homeostasis, transport, and signal pathways (Chen et al., 2018; Huang et al., 2021; Zhang et al., 2023; Xian et al., 2026). Mutation of auxin efflux transporter encoding gene PIN-FORMED 1b (*OsPIN1b*) (Wang et al., 2022), or overexpression of *indole-3-acetic acid glucosyltransferase* (*OsIAAGLU*) (Yu et al., 2019) and *OsGH3* (Zhao et al., 2013; Zhang et al., 2015), which encode indole-3-acetic acid glucosyltransferase and IAA-amido synthetase, respectively, all result in reduced free auxin content in lamina joint and subsequently an enlarged leaf angle. Gain-of-function mutation of *LEAF INCLINATION1* (*LC1*)/*OsGH3-1* shows exaggerated FLA (Zhao et al., 2013). Overexpression of *OsmiR393* or suppression of its silencing targets *OsTIR1* and Auxin Signaling F-box 2 (*OsAFB2*) results in increased leaf angle (Bian et al., 2012). The SPOC domain-containing transcriptional repressor LEAF INCLINATION3 (OsLC3) and OsLC3-Interacting Protein 1 (OsLIP1) synergistically inhibit the expression of *OsGH3*.*2* and *OsIAA12* (Chen et al., 2018). OsGH3.2 reduces free auxin levels, while OsIAA12 suppresses the auxin signaling cascade through Auxin Response Factor17 (OsARF17) (Chen et al., 2018). Deficiency of *OsLC3* or *OsARF17*, or overexpression of *OsIAA12*, all lead to an exaggerated leaf angle (Chen et al., 2018). OsARF6 and OsARF17 directly activate the expression of the *MAPKKK* family member *Increased Leaf Angle1* (*ILA1*), control FLA by modulating the secondary cell wall levels of sclerenchyma cells in lamina joint (Huang et al., 2021).

BRs have prominent effects on FLA control in rice, as increasing BR levels or enhancing BR signaling promotes leaf inclination (Liu et al., 2024). BR biosynthesis gene deficient mutants of rice, including *dwarf4, dwarf11* (*d11*), *ebisu dwarf* (*d2*), *BR-deficient dwarf1* (*brd1*), and *brd2*, show erect leaves (Hong et al., 2002; Hong et al., 2003; Hong et al., 2005; Tanabe et al., 2005; Sakamoto et al., 2006). Suppression or knockout of the positive components in the BR signaling, such as *Brassinosteroid Upregulated1* (*OsBU1*) (Tanaka et al., 2009), *Increased Lamina Joint Inclination1* (*OsILI1*) (Zhang et al., 2009), *Brassinosteroid Insensitive1* (*OsBRI1*) (Yamamuro et al., 2000), *DWARF AND LOW-TILLERING* (*OsDLT*) (Tong et al., 2012), Brassinosteroid Insensitive 1-associated kinase 1 (*OsBAK1*) (Li et al., 2009a), Reduced Leaf Angle1 (*RLA1*)/ *SMALL ORGAN SIZE1* (*SMOS1*) (Qiao et al., 2017), *OsMED25* (Ren et al., 2020), *BRASSINAZOLE-RESISTANT1* (*OsBZR1*) (Bai et al., 2007; Yu et al., 2021) and *OsBZR2* (Liu et al., 2021), all results in erect leaves. Various factors are involved in leaf angle control associated with BR-mediated progress, such as transcriptional regulator OsGATA7, CYC U4;1 and *Put On Weight 1* (*POW1*) (Sun et al., 2015; Zhang et al., 2018; Zhang et al., 2021). Knockout mutant and gain-of-function mutant of *OsBZR1* show erect and droopy leaf angle, respectively (Qiao et al., 2017; Yu et al., 2021). As a conserved transcription factor acts downstream of Glycogen Synthase Kinases (OsGSKs), OsBZR1 mediates BR signal to leaf angle by regulating various genes (Liu et al., 2024). For example, a pair of antagonistic HLH/bHLH factors OsILI1 and ILI1 binding bHLH (IBH1) (Zhang et al., 2009), the glutamate receptor-like channel protein OsGLR3.4 (Yu et al., 2021), and the R2R3-MYB transcription factor FOUR LIPS (OsFLP) (Liu et al., 2024). *GLUTAMATE RECEPTOR-LIKE3*.*4* (*OsGLR3*.*4*) mutant exhibits reduced leaf angle mainly due to inhibited adaxial cell elongation (Yu et al., 2021). OsFLP mediates BR signals to lignin deposition in vascular bundles and sclerenchyma cells within lamina joint, and feedback regulates *OsGSK1* transcription and OsBZR1 phosphorylation status (Liu et al., 2024). In the process of regulating leaf angle, the CCCH-type zinc finger protein OsLIC serves as an antagonistic transcription factor of OsBZR1 (Qu et al., 2012), the mediator subunit OsMED25 functions as a corepressor of OsBZR1 (Ren et al., 2020), and the APETALA2 family transcription factor RLA1/SMOS1 can enhance the transcriptional activity of OsBZR1, and is a phosphorylates substrate of OsGSK2 (Qiao et al., 2017).

As mentioned above, the magnitude of rice leaf angle exhibits a positive correlation with BR levels and a negatively correlation with auxin content in the lamina joint (Huang et al., 2021). However, the molecular mechanisms underlying their antagonistic interaction in leaf angle regulation remained elusive. In this study, we demonstrate that BR signaling represses OsYUCCA8 (OsYUC8)-mediated auxin biosynthesis to control FLA by regulating SCW formation. Genetic evidence shows that *OsYUC8* deficiency exhibits an increased FLA accompanied by disrupted SCW deposition. Moreover, OsBZR1, a key transcription factor in BR signaling, directly binds to *OsYUC8* promoter and represses its expression, thereby decreases auxin-mediated SCW biosynthesis. Finally, field trials further reveal that OsBZR1 is important for improving yield performance under dense planting conditions. Together, our study not only uncovers an antagonistic BR-auxin interaction that governs FLA but also establishes a genetic strategy for manipulating crop architecture to maximize yield under dense planting regimes.

## Results

### Antagonistic hormone pathways control rice FLA

FLA is regulated by two well-known hormones, auxin and BR (Cao et al., 2022). To study the role of these two hormones in the genetic regulation of FLA in rice, we examined a panel of mutants impaired in auxin or BR signaling and biosynthesis (Fig. 1A). All six auxin-related mutants examined (*mao huzi 10* [*mhz10*], *osafb2, osarf6*/*12*/*17, ospin1b*) exhibited consistently increased FLA compared with wild-type (WT) plants (Bian et al., 2012; Huang et al., 2021; Zhou et al., 2022; Zhang et al., 2023). In striking contrast, each of the six BR-related mutants (*d2, brd2, osdwarf4, osbri1, osbzr1, osbak1*) displayed a markedly decreased FLA (Yamamuro et al., 2000; Hong et al., 2003; Sakamoto et al., 2006; Bai et al., 2007; Li et al., 2009b). These opposing phenotypes suggest that auxin and BR pathways exert antagonistic effects on leaf inclination.

**Fig. 1.**
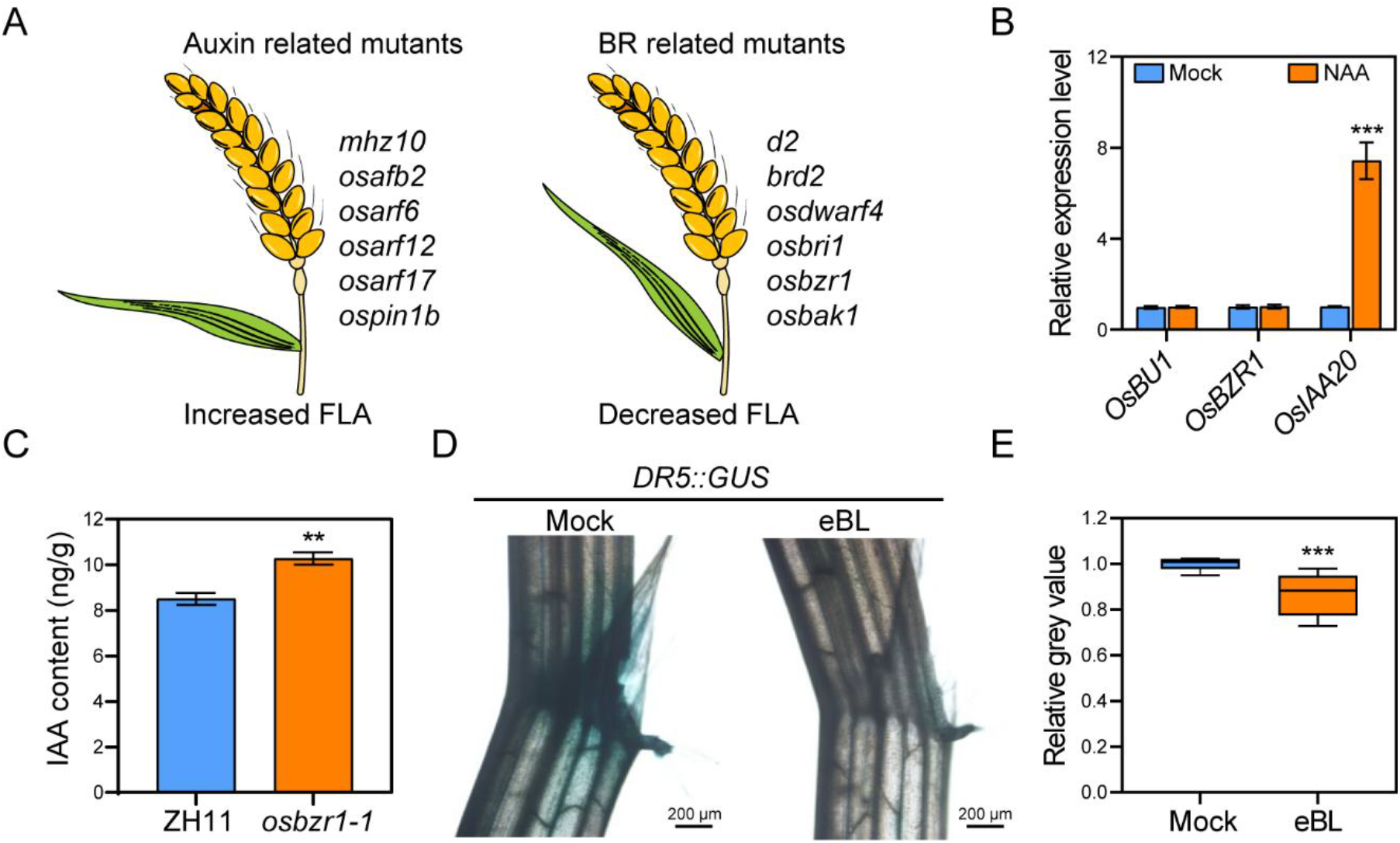
BR-auxin antagonism regulates FLA. **A)** Auxin and BR mutants exhibit antagonistic FLA phenotypes. **B)** The expression of BR signaling genes is unaffected by NAA treatment in the lamina joints of flag leaf of 100-day-old plants. Error bars are ± SE, n = 4 biological replicates. Student’s *t*-test: \*\*\**P* < 0.001. **C)** Auxin levels are increased in the lamina joints of 100-day-old *osbzr1-1* mutants. Error bars are ± SE, n = 3 biological replicates. Student’s *t*-test: \*\**P* < 0.01. **D)** GUS staining of *DR5::GUS* transgene plants treated with/without 10 μM eBL. Scale bars, 200 μm. **E)** Statistical graph of the relative grey value from **(D)**. Student’s *t*-test: \*\*\**P* < 0.001, n = 10 biological replicates.

To investigate the relationship between these two hormones, we treated the lamina joint of the flag leaf with 1-naphthaleneacetic acid (NAA, a synthetic auxin analog) and 24-epibrassinolide (eBL, bioactive BR). Exogenous NAA (100 nM) treatment did not affect the expression of BR-related genes (Fig. 1B), in contrast, auxin levels were elevated in the *osbzr1* mutants (Fig. 1C). Consistent with those observations, eBL (10 μM) treatment suppressed the signal intensity of the auxin reporter DR5::GUS (Fig. 1, D and E). These results indicate that BR regulates FLA by negatively modulating auxin biosynthesis. Collectively, these genetic and cellular results establish that proper FLA requires a balance between auxin and BR signaling and provide a foundation for dissecting the molecular mechanisms by which these two hormone pathways coordinately regulate leaf architecture.

### OsYUC8 regulates FLA via SCW deposition in sclerenchyma cells

*YUC* (flavin monooxygenase-like) gene families encode the key rate-limiting enzymes in the indole-3-pyruvic acid (IPyA) pathway, which represents the primary auxin biosynthesis route in plants (Zhao, 2010). Among the *YUC* genes highly expressed in the lamina joint of rice flag leaf, eBL treatment assays revealed that only *OsYUC8* was significantly repressed by exogenous eBL (Supplementary Fig. S1). Intriguingly, mutants of *OsYUC8* consistently displayed loose plant architecture under field conditions (Fig. 2A). Detailed morphological analysis revealed that the architectural defects of *osyuc8-1* and *osyuc8-2* mutants were specially resulted from significantly increased FLA (Fig. 2, A to C) (Kong et al., 2024b). These results indicate that OsYUC8 plays a major role in the regulation of FLA.

**Fig. 2.**
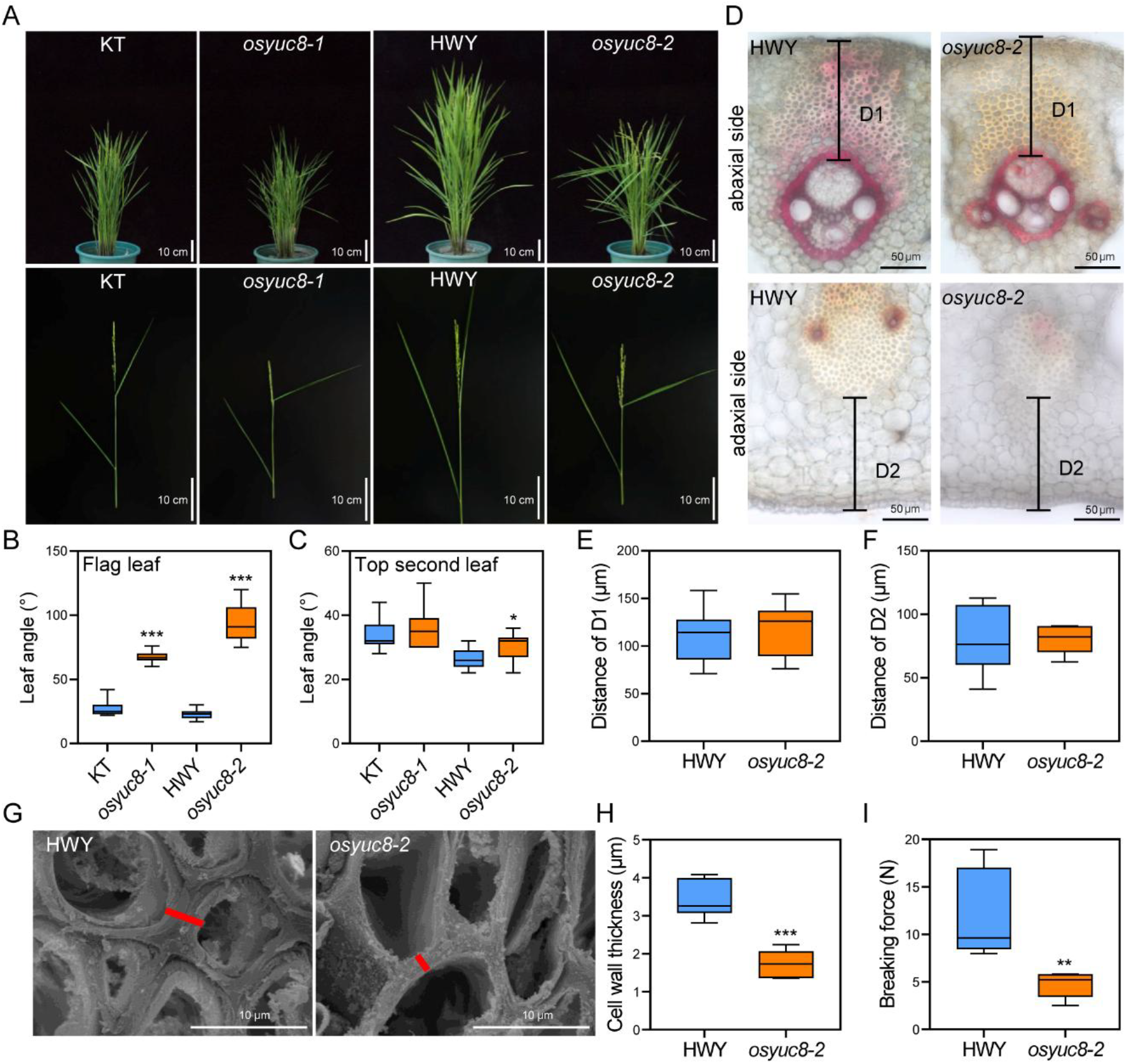
*OsYUC8*-mediated auxin biosynthesis governs FLA. **A)** The representative images of 100-day-old WT and *OsYUC8* mutants. Scale bars, 10 cm. **B and C)** Analysis of the FLA **(B)** and the top second leaf angle **(C)** in WT and *OsYUC8* mutants. Student’s *t*-test: \**P* < 0.05, \*\*\**P* < 0.001, n = 15 biological replicates. **D)** Cross sections from flag leaf lamina joints at the ripening stage after staining with phloroglucinol-HCl. D1 designates the region interposed between the adaxial epidermis and the sclerenchyma bundles. D2 represents the distance of lamina joint adaxial margin to vascular bundles. Scale bars, 50 μm. **E and F)** The distance of D1 **(E)** and D2 **(F)**. n ≥ 10 biological replicates. **G)** Representative cell wall SEM images of abaxial sclerenchyma tissue in the lamina joints of HWY and *osyuc8-2*. Scale bars, 10 μm. Red lines indicate the measured regions. **H)** Statistical graph of the cell wall thickness from **(G)**. Student’s *t*-test: \*\*\**P* < 0.001, n = 5 biological replicates. **I)** Mechanical measurement showing breaking force between HWY and *osyuc8-2*. Student’s *t*-test: \*\**P* < 0.01, n = 5 biological replicates.

To elucidate the cellular mechanisms underlying the observed phenotypes, we performed morphological and biochemical analysis. Typically, leaf angle is mainly determined by the pushing force of adaxial parenchyma cells and the mechanical strength of vascular bundles and sclerenchyma tissues within lamina joint(Zhou et al., 2017; Cao et al., 2022). Our comprehensive morphometric analysis demonstrated that *OsYUC8* deficiency did not alter fundamental lamina joint architecture given that the lengths of abaxial and adaxial sides of the lamina joint in *osyuc8-1* and *osyuc8-2* were similar with those of WT plants (Supplementary Fig. S2), and detailed cytological observation showed that the lengths of D1 and D2 in lamina joint of *osyuc8-2* were also normal in comparison with that of WT plants (Fig. 2, D to F). However, biochemical staining and scanning electron microscopy (SEM) imaging, and cell wall component analysis indicated that the lignin and cellulose content, as well as cell wall thickness, in the abaxial sclerenchyma cells of *osyuc8-2* were significantly reduced compared with those of WT plants (Fig. 2, D, G and H, Supplementary Fig. S3). Consistent with these findings, dynamic mechanical analysis showed that the breaking force at the lamina joint of *osyuc8-2* was significantly smaller than that of WT lines (Fig. 2I), indicating a marked reduction in the mechanical strength of lamina joint in the *OsYUC8* mutants. Together, these results demonstrate that OsYUC8 regulates FLA through a mechanism primarily involving SCW deposition in abaxial sclerenchyma cells, rather than through gross morphological alterations of the lamina joint.

### OsBZR1 directly represses *OsYUC8* expression

To delineate the regulatory network controlling OsYUC8-mediated FLA determination, we performed systematic molecular screening and validation. Yeast one-hybrid screening of a rice cDNA library using a 3.0 kb *OsYUC8* promoter fragment identified ten putative transcriptional regulators (Fig. 3A). Among these, OsBZR1, the central transcription factor in BR signaling, emerged as the most promising candidate based on strong interaction signal intensity (Fig. 3B) and known functional relevance to leaf angle regulation (Bai et al., 2007). The promoter sequence of *OsYUC8* contains eight E-BOX and one BRRE elements, which are supportive for the binding of OsBZR1 (Fig. 3C). Furthermore, chromatin immunoprecipitation (ChIP)-qPCR confirmed specific binding of OsBZR1 to two regions: CF1 (-2492 to -2366 bp) and CF2 (-2284 to -2175 bp) (Fig. 3, C and D, Supplementary Fig. S4). These results suggest that BR acts upstream of auxin in regulating FLA.

**Fig. 3.**
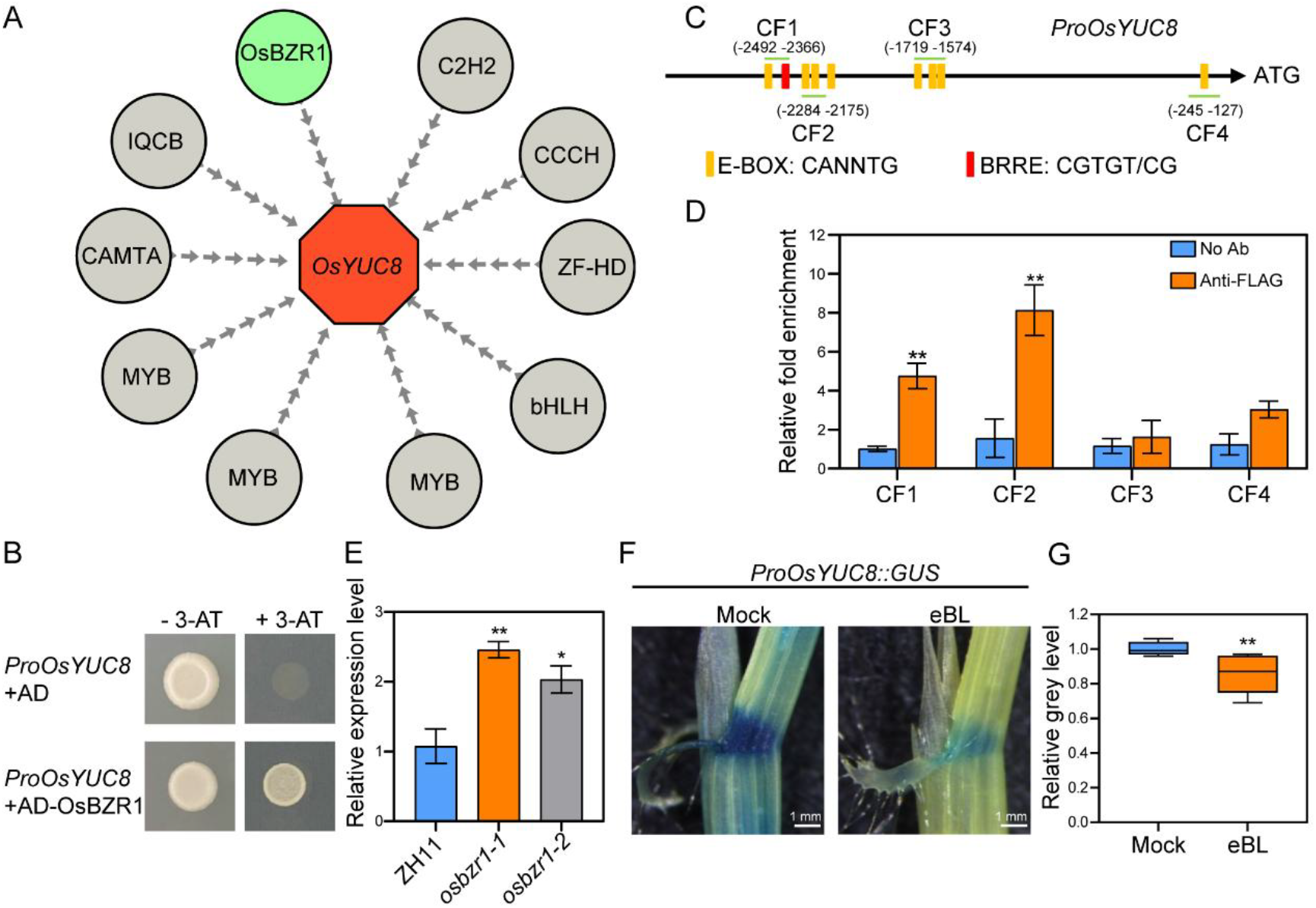
OsBZR1 directly represses *OsYUC8* expression. **A)** Yeast one-hybrid screening identifies putative regulators of *OsYUC8*. **B)** Yeast one-hybrid confirms OsBZR1 binding to the *OsYUC8* promoter. The experimental group was cultured on SD medium lacking leucine and histidine supplemented with 3 mM 3-AT. **C)** Diagram of the *OsYUC8* promoter region. Orange box: E-BOX (CANNTG); red box: BRRE (CGTGT/CG). **D)** ChIP–qPCR assays to detect interaction between OsBZR1 and CF1/2 within the *OsYUC8* promoter in lamina joints. CF1 fragment contains one E-BOX and one BRRE element; CF2 and CF3 comprise two and three E-BOX elements, respectively; CF4 contains one E-BOX element. Error bars indicate ± SE from 3 biological replicates. Student’s *t*-test: \*\**P* < 0.01. **E)** RT-qPCR showed relative expression level of *OsYUC8* in ZH11, *osbzr1-1* and *osbzr1-2* mutants. Error bars indicate ± SE from 3 biological replicates. Student’s *t*-test: \**P* < 0.05, \*\**P* < 0.01. **F)** GUS staining of *ProOsYUC8::GUS* transgene plants treated with/without 10 μM eBL. Scale bars, 1 mm. **G)** Statistical graph of the relative grey value from **(F)**. Student’s *t*-test: \*\**P* < 0.01, n = 10 biological replicates.

Therefore, it is crucial to clarify the role of OsBZR1 binding to the *OsYUC8* promoter in regulating its expression. RT-qPCR showed that the expression of *OsYUC8* was significantly upregulated in *osbzr1-1* and *osbzr1-2* mutants (Fig. 3E), indicating the inhibitory effect of OsBZR1 on *OsYUC8* expression. External eBL treatment of *ProOsYUC8::GUS* transgenic plants showed reduced GUS activity (Fig. 3, F and G). These results indicate that OsBZR1 negatively modulates FLA via directly repressing *OsYUC8* expression in the lamina joint.

### OsBZR1 genetically acts upstream of *OsYUC8*

To verify the roles of OsBZR1 in regulating FLA, we characterize the phenotype of *OsBZR1* mutants. Consistent with previous studies that constitutively activated BR signaling of plants exhibit a large leaf angle (Bai et al., 2007), *OsBZR1* mutants exhibited smaller FLA (Fig. 4, A to C). Similar to that of the *OsYUC8* mutants, the lengths of D1 and D2 in the lamina joints of *osbzr1-1* and *osbzr1-2* showed no significant difference from those of WT lines (Fig. 4, D to F). Phloroglucinol staining and SEM imaging results revealed that the lignin content and thickness of the abaxial sclerenchyma cell wall in *osbzr1* mutants were significantly increased compared with the WT plants (Fig. 4, D, G and H). Consistently, lignin and cellulose content was elevated in *osbzr1* mutants (Supplementary Fig. S5). Correspondingly, the breaking force at the lamina joint of both *osbzr1-1* and *osbzr1-2* was significantly higher than that of WT plants (Fig. 4I). These results indicate that OsBZR1 regulates FLA by suppressing SCW biosynthesis.

**Fig. 4.**
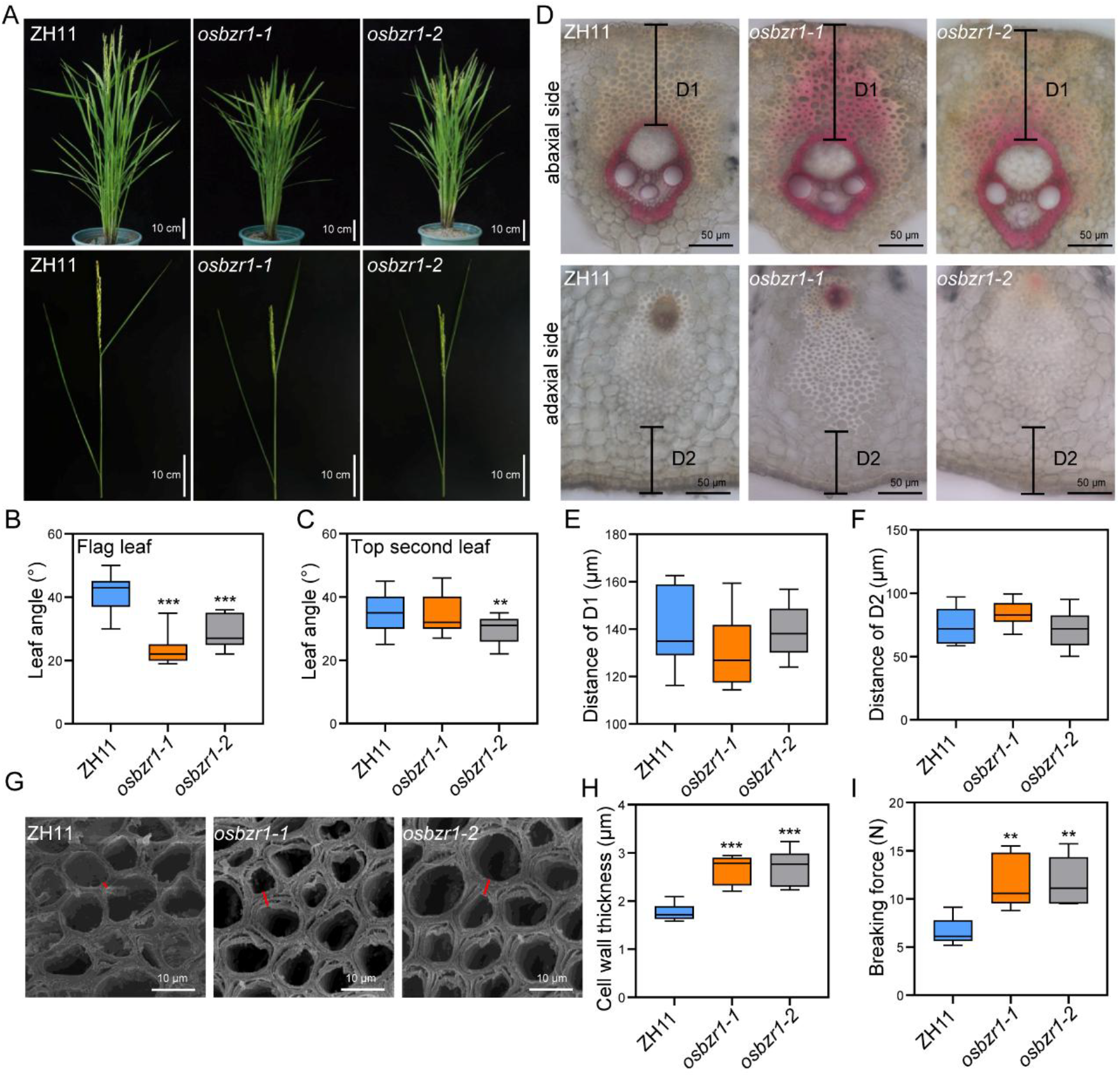
Mutations of *OsBZR1* lead to thickened cell wall in the lamina joint of flag leaf. **A)** The representative images of 100-day-old WT and *OsBZR1* mutants. Scale bars, 10 cm. **B and C)** Analysis of the flag leaf **(B)** and the top second leaf angle **(C)** in WT and *OsBZR1* mutants. Student’s *t*-test: \*\**P* < 0.01, \*\*\**P* < 0.001, n = 15 biological replicates. **D)** Cross sections from flag leaf lamina joints at the ripening stage after staining with phloroglucinol-HCl. D1 designates the region interposed between the adaxial epidermis and the sclerenchyma bundles. D2 represents the distance of lamina joint adaxial margin to vascular bundles. Scale bars, 50 μm. **E and F)** The distance of D1 **(E)** and D2 **(F)**. n ≥ 10 biological replicates. **G)** Representative cell wall SEM images of abaxial sclerenchyma tissue in the flag leaf lamina joints of ZH11, *osbzr1-1* and *osbzr1-2* mutants. Scale bars, 10 μm. Red lines indicate the measured regions. **H)** Statistical graph of the cell wall thickness from **(G)**. Student’s *t*-test: \*\*\**P* < 0.001, n = 6 biological replicates. **I)** Dynamic mechanical analysis showing breaking force of ZH11, *osbzr1-1* and *osbzr1-2* mutants. Student’s *t*-test: \*\**P* < 0.01, n = 5 biological replicates.

To further dissect the genetic relationship between OsBZR1 and OsYUC8, we generated the *osbzr1 osyuc8* double mutant (*dm*) by knocking out *OsYUC8* in the *osbzr1* background (Supplementary Fig. S6). Clearly, the *dm* line exhibited a larger FLA, comparable to that of WT plants, and similar to the phenotype observed in the *OsYUC8* mutants (Supplementary Fig. S7). Moreover, the lengths of D1 and D2 in the lamina joints of *dm* showed no significant difference from those of *OsYUC8* mutants (Supplementary Fig. S8). Collectively, these results indicate that OsBZR1 acts upstream of OsYUC8 to regulate rice FLA through modulating SCW formation.

### *OsBZR1* mutants perform well in high dense planting conditions

A more upright leaf angle can effectively mitigate the detrimental side effects associated with dense planting systems (Sinclair and Sheehy, 1999). To further evaluate the contribution of FLA to grain yield under varying planting densities, we grew the *osyuc8* and *osbzr1* mutants under dense planting conditions, and systematically characterized their yield-related agronomic traits. Compared with WT plants, the *osyuc8-2* mutants exhibited a substantial reduction in grain yield under high-density planting conditions, reflected in a significantly increased percentage of empty grains, along with a moderate negative effect on grain size (Fig. 5, A to F, Supplementary Fig. S9). In contrast, high-density planting exerted a much less inhibitory effect on grain yield in the *osbzr1-1* mutants relative to the WT plants (Fig. 5, G to L, Supplementary Fig. S9). This mutant maintained a markedly higher seed-setting rate without compromising grain size, which is a key determinant of final yield (Fig. 5, G to L, Supplementary Fig. S9). Taken together, these results indicate that OsBZR1 represents a promising target for achieving a more upright leaf angle while sustaining yield under high-density cultivation conditions.

**Fig. 5.**
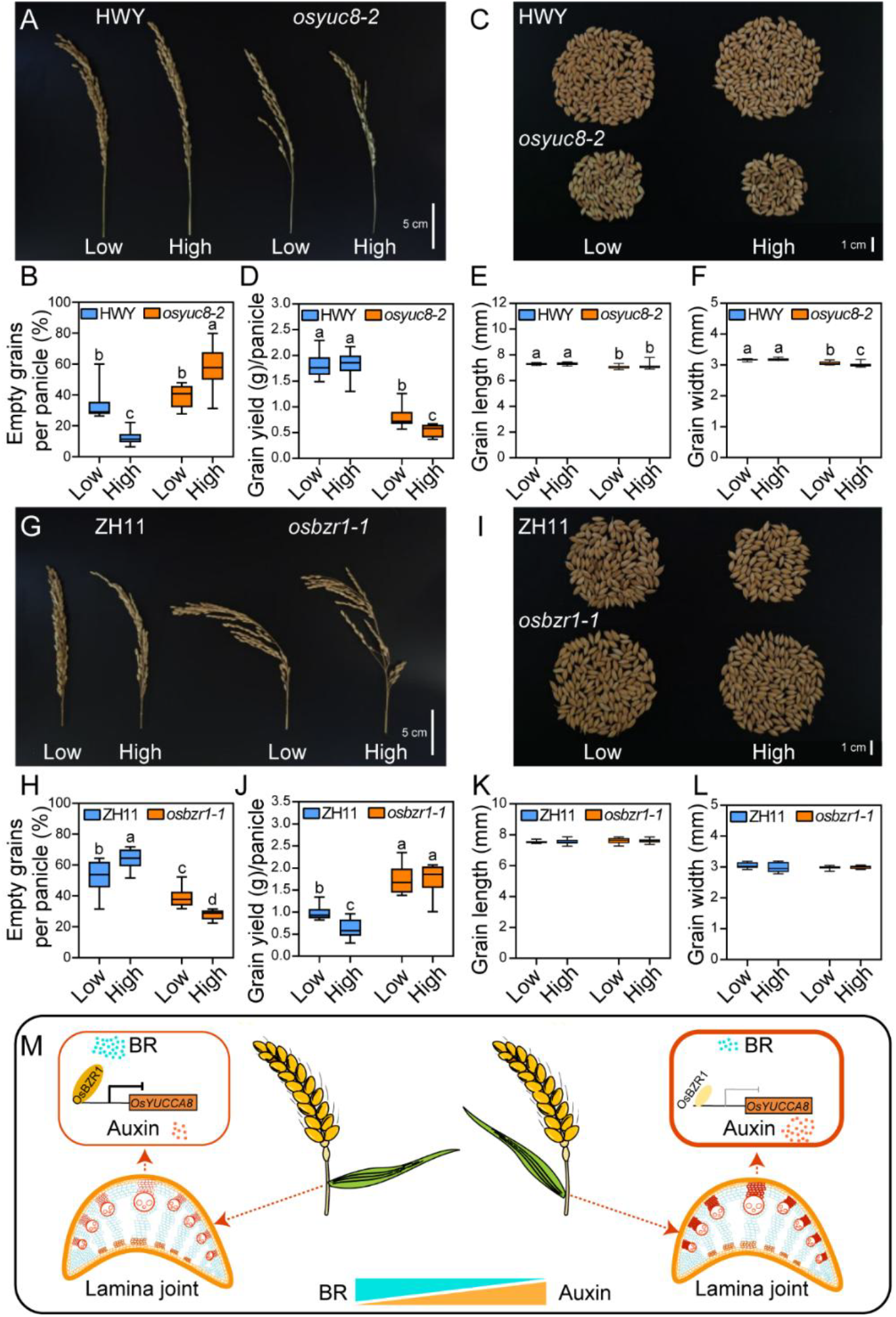
OsBZR1 confers yield advantage in rice under high plant density. **A)** Panicle phenotypes of WT and *osyuc8-2* mutants under different planting densities. Scale bars, 5 cm. **B)** The percentage of empty grain per panicle of WT and *osyuc8-2* mutants under different planting densities: low (25 plants m^−2^), and high (49 plants m^−2^). Different letters indicate significant differences, *P* < 0.01 by one-way ANOVA analysis, n ≥ 18 biological replicates. **C)** Plump grains from three panicles of WT and *osyuc8-2* mutants under different planting densities. Scale bars, 1 cm. **D)** Grain yield per panicle of WT and *osyuc8-2* mutants under different planting densities. Different letters indicate significant differences, *P* < 0.01 by one-way ANOVA analysis, n = 19 biological replicates. **E)** Grain length of WT and *osyuc8-2* mutants under different planting densities. Different letters indicate significant differences, *P* < 0.01 by one-way ANOVA analysis, n = 18 biological replicates. **F)** Grain width of WT and *osyuc8-2* mutants under different planting densities. Different letters indicate significant differences, *P* < 0.01 by one-way ANOVA analysis, n = 19 biological replicates. **G)** Panicle phenotypes of WT and *osbzr1-1* mutants under different planting densities. Scale bars, 5 cm. **H)** The percentage of empty grain per panicle of WT and *osbzr1-1* mutants under different planting densities. Different letters indicate significant differences, *P* < 0.01 by one-way ANOVA analysis, n = 15 biological replicates. **I)** Plump grains from three panicles of WT and *osbzr1-1* mutants under different planting densities. Scale bars, 1 cm. **J)** Grain yield per panicle of WT and *osbzr1-1* mutants under different planting densities. Different letters indicate significant differences, *P* < 0.01 by one-way ANOVA analysis, n = 15 biological replicates. **K)** Grain length of WT and *osbzr1-1* mutants under different planting densities. n = 15 biological replicates. **L)** Grain width of WT and *osbzr1-1* mutants under different planting densities. n = 15 biological replicates. **M)** BR and auxin antagonistically regulate rice flag leaf angle. High BR levels activate OsBZR1, which represses the auxin biosynthetic gene *OsYUC8*, leading to low auxin accumulation. This results in thinner cell walls in the lamina joint sclerenchyma, providing weak mechanical support and thus a larger FLA. Conversely, low BR levels relieve *OsYUC8* repression, increasing auxin accumulation, which promotes thicker cell walls, strong mechanical support, and a smaller FLA.

This study elucidates an unreported BR-auxin signaling cascade governing rice FLA regulation that BR signaling acts upstream of local auxin biosynthesis through direct binding of OsBZR1 to the *OsYUC8* promoter and transcriptional repression of *OsYUC8*. The OsBZR1-OsYUC8 module controls secondary cell wall deposition in abaxial sclerenchyma and thus lamina joint mechanical strength. These findings fundamentally advance our understanding of phytohormone crosstalk in plant architecture regulation, providing both theoretical insights and practical tools for crop improvement.

## Discussion

Cereal crop architecture, particularly FLA, is a key determinant of canopy structure and yield potential under high-density cultivation (Sinclair and Sheehy, 1999). Although the individual roles of BR signaling and auxin biosynthesis in plant development are well established (Sakamoto et al., 2006; Huang et al., 2021), the mechanistic link between these hormonal pathways and the biomechanical properties of the leaf lamina joint has been a long-standing question. Here, we elucidate a molecular framework in which BR signaling governs FLA by directly repressing auxin biosynthesis at the transcriptional level, specifically through the gene *OsYUC8*, thereby modulating SCW formation, a process critical for erect flag leaf architecture (Fig. 5M). This finding reframes our understanding of hormone action from a model of parallel pathways to one of hierarchical, antagonistic regulation, with BR acting as a master brake on auxin production to achieve a specific developmental outcome.

At the core of this regulatory module is the direct interaction between a master transcription factor (OsBZR1) and a rate-limiting biosynthetic enzyme (OsYUC8) (Bai et al., 2007; Qin et al., 2017; Kong et al., 2024a). We identify OsBZR1 as a direct repressor of *OsYUC8* transcription (Fig. 3), providing a mechanistic explanation for the long-observed antagonism between BR and auxin. Genetic analyses demonstrate that loss-of-function mutations in *OsYUC8* recapitulate the defective SCW formation and increased FLA phenotypes (Fig. 2) observed in both BR-hyperresponsive mutants and auxin signaling impaired lines(Bai et al., 2007; Tanaka et al., 2009; Huang et al., 2021), indicating that BR signaling primarily modulates FLA by fine-tuning the auxin pool to control SCW biosynthesis. Our study reveals that OsYUC8-mediated local auxin biosynthesis promotes SCW deposition, while BR signaling antagonizes this process by suppressing *OsYUC8* expression, thereby fine-tuning SCW biosynthesis in specific lamina joint tissues (Fig. 2 to 4). The resulting decrease in mechanical rigidity, likely on the abaxial side, enables a larger FLA. This mechanistic insight resolves a key developmental question: how plants achieve precise, localized tissue modifications for organ angle control without globally disrupting growth, through cell-type-specific transcriptional regulation of a core metabolic pathway.

From an agricultural biotechnology perspective, these findings provide a targeted solution to the long-standing density-yield trade-off. The *osbzr1* mutants exemplify precision breeding: by disrupting a negative regulator in this BR-auxin-SCW module, we achieve elevated auxin-mediated SCW synthesis and the desired erect leaf phenotype without the pleiotropic dwarfism or reproductive defects associated with broader hormonal perturbations (Fig. 4). Field evaluations confirm improved yield under high-density cultivation (Fig. 5), demonstrating the agronomic viability of this approach. By targeting a tissue-specific trait rather than systemic growth, this strategy preserves overall biomass and grain production, providing a molecular toolkit for designing ideal plant architecture suited to mechanized, high-density agriculture. In conclusion, this work not only deciphers a fundamental antagonistic hormonal interaction but also provides a precise blueprint for engineering crop architecture to meet the sustainability and productivity demands of modern agriculture.

## Materials and Methods

### Plant materials and growth conditions

The genetic backgrounds of *osbzr1-1, osbzr1-2*, and *osbzr1-1 osyuc8-3* in this study are Zhonghua11 (ZH11) (*Oryza sativa, japonica*). The backgrounds of *osyuc8-1/rein7-1, osyuc8-2/oscow1-2/rein7-2* are Kitaake (KT) (*Oryza sativa, japonica*), Hwayoung (HWY) (*Oryza sativa, japonica*), respectively. *Osyuc8-1* harbors a single base pair substitution in the fourth exon resulting in a stop codon of *OsYUC8* gene. *Osyuc8-2* is a T-DNA insertion mutant at the second intron of *OsYUC8* gene. Plants were cultured in a paddy field at Sanya (18º N, 109º E) in the winter and Shanghai (30º N, 121º E) in the summer. Rice transgenic transformation was conducted by EDGENEBIOT Company (https://www.edgene.com.cn/). In the field trials, two planting densities were established for yield evaluation: a high-density treatment of 49 plants m^−2^ and a low-density treatment of 25 plants m^−2^.

### Lamina joint slicing and microscopic imaging

Lamina joints were harvested and embedded in 5% (w/v) agarose. Serial sections (50 ± 5 μm) were prepared using a precision vibrating microtome (Leica, VT1200S). Tissue sections were subsequently stained with acidified phloroglucinol solution (2% w/v in 95% ethanol, HCl-activated) for 5 min at 25°C prior to bright-field microscopy analysis. Measurements of cell length on the adaxial and abaxial surfaces were performed utilizing ImageJ software.

### Hormone treatment

Seeds of *ProYUC8::GUS* and *DR5::GUS* transgenic lines were germinated in water for about 4 days, and transplanted to 96-well boxes containing 1 L Yoshida solution. After 10 days of growth, leaf joints were sprayed with 10 μM eBL, and double distilled water was conducted as negative control. After treatment for 6 h, lamina joint materials were collected for subsequent experiments.

### RNA isolation and RT-qPCR

RNA extractions were performed using the TRNzol Universal RNA Reagent (TIANGEN, DP424) according to the manufacturer’s protocol. Then complementary DNA (cDNA) was synthesized using ToloScript ALL-in-one RT EasyMix (TOLOBIO, 22107). qPCR was carried out using 2×Q5 SYBR qPCR Master Mix (Universal) (TOLOBIO, 22208) on a Light-Cycler® instrument (Roche, Germany). Rice *UBIQUITIN* (*Os05g06770*) was used as internal reference gene, all experiments comprise three biological replicates and three technical replicates. RT-qPCR primers are listed in *SI Appendix*, Table S1.

### Measurements of auxin content

Lamina joints of WT and mutant plants were excised in the field, immediately frozen in liquid nitrogen, and stored at —80°C. Subsequently, the samples were sent to Metware Company (https://www.metware.cn/) for auxin level determination.

### Measurements of lignin and cellulose content

For lignin content determination, ground samples in liquid nitrogen were washed twice with 95% (v/v) ethanol, subjected to centrifugation at 700g for 2 min at 25°C, and then oven-dried at 30°C to constant weight. Aliquots of 20 mg of the dried material were transferred to screw-cap centrifuge tubes containing 2 mL of 25% (v/v) acetyl bromide in glacial acetic acid and incubated at 70°C for 30 min. Following complete digestion, a 0.2-mL portion of the digest was combined with 0.45 mL of 2 M NaOH, 0.05 mL of 7.5 M hydroxylamine-HCl, and 3.3 mL of glacial acetic acid. The resulting mixture was diluted to a final volume of 10 mL with glacial acetic acid, and the absorbance of the supernatant was measured at 280 nm.

For cellulose content determination, lamina joints were dried and reduced to fine powders using a ball mill. Alcohol-insoluble residues were prepared, destarched, and then hydrolyzed with 2 M trifluoroacetic acid. Subsequently, the residues were treated with updegraff reagent. The resulting pellets were subjected to 72% (v/v) sulfuric acid, after which cellulose content was quantified using the anthrone method.

### GUS staining

Rice materials were harvested and vacuum-infiltrated with GUS staining solution (50 mM NaHPO_4_ pH 7.2, 0.5% Triton X-100, 1 mM X-Gluc) for 30 minutes and subsequently incubated at 37°C for 12 hours in the dark, and 70% ethanol were used for decolorize.

### Scanning electron microscopy imaging

The flag leaf lamina joints of WT, *OsYUC8*, and *OsBZR1* mutants were collected and fixed in FAA solution (formaldehyde : acetic acid : ethanol : water = 5 : 5 : 63 : 27) for 12 hours. Then, the samples were dehydrated in an ethanol gradient series (70%, 80%, 90%, 95% and 100% ethanol), followed by critical point drying (Leica EM CPD300). Samples were stick to a copper platform and gold-coated using a vacuum coater (Leica EM SCD050). Finally, samples were observed and photographed by SEM (Hitachi S-3400N II).

### Mechanical force measurement

Sections approximately 2 cm above and below the flag leaf lamina joints were excised. Segments were then clamped at both extremities within a dynamic mechanical analyzer (TAINSTRUMENTS, Q850) and subjected to continuously increasing tensile force until rupture occurred. The minimum force required to induce fracture was recorded.

### Yeast one-hybrid assay

For yeast one-hybrid screens, the 3000 bp promoter sequences of *OsYUC8* were cloned to pHis vector. The constructs were co-transformed with rice transcription factor cDNA library into AH109. Following incubation at 30°C for 72 hours, individual colonies were isolated for subsequent validation assays. Yeast one-hybrid library screen was conducted by Oebiotech Company (https://www.oebiotech.com/).

### ChIP-qPCR

OsBZR1-FLAG transgenic seedlings were harvested and crosslinked in 1% (v/v) formaldehyde. Subsequently, tissues were homogenized in liquid nitrogen followed by chromatin extraction via nuclei lysis. The solubilized chromatin was subjected to sonication for DNA fragmentation. ChIP was then performed using anti-FLAG antibody (1:200 dilution; Abclonal, AE092) conjugated to protein G magnetic beads (Thermo, 01134323). Precipitated DNA was purified and analyzed by quantitative PCR (qPCR). Protein G coupled beads without antibody served as negative controls.

### Western bolt

Total protein extracts were isolated from transgenic plants and immunoblotted with mouse anti-FLAG antibody (1:2000 dilution; Abclonal, AE092). Histone H3 rabbit antibody (1:1000 dilution; Abclonal, A22348) served as the internal loading control. Following extensive washes, membranes were probed separately with horseradish peroxidase (HRP)-conjugated goat anti-mouse (1:5000 dilution; Abmart, M21001) and goat anti-rabbit (1:5000 dilution; Abmart, M21002) secondary antibodies. Protein-antibody complexes were subsequently visualized using enhanced chemiluminescence (Share bio, SB-WB012) with exposure times optimized for linear signal detection.

### Accession numbers

Sequence data from this article can be found in the GenBank/EMBL data libraries under the following accession numbers: *OsYUC8* (*Os03g0162000*), *OsBZR1* (*Os07g0580500*), *OsBU1* (*Os06g0226500*), *OsIAA20* (*Os06g0166500*), *OsYUC1* (*Os01g0645400*), *OsYUC3* (*Os01g0732700*), *OsYUC5* (*Os12g0512000*), *OsYUC6* (*Os07g0437000*), *OsYUC7* (*Os04g0128900*), *OsYUC10* (*Os01g0274100*).

## Supplemental Data

**Supplemental Figure S1**. *OsYUC8* expression is inhibited by external eBL treatment.

**Supplemental Figure S2**. The lamina joint morphology of flag leaf is normal in *osyuc8* mutants.

**Supplemental Figure S3**. *Osyuc8-2* mutants exhibit decreased lignin and cellulose levels.

**Supplemental Figure S4**. Validation of OsBZR1-FLAG transgenic lines by western blot.

**Supplemental Figure S5**. Increased lignin and cellulose content in *osbzr1-1* mutants.

**Supplemental Figure S6**. Knockout *OsYUC8* in *osbzr1-1* background.

**Supplemental Figure S7**. Genetic epistasis of *OsYUC8* and *OsBZR1* in controlling FLA.

**Supplemental Figure S8**. Lamina joint morphology of flag leaf of *osbzr1-1 osyuc8-3* mutant is normal.

**Supplemental Figure S9**. Grain size of WT, *osyuc8-2*, and *osbzr1-1* mutants under different planting densities.

**Supplemental Table S1**. Primers used in this study

## Author Contributions

X. K., and G. H. designed research; S. Y., Q. L., Y. X., D. T., Y.Y., and Z.Q. performed research;

S. Y., and X. K. analyzed data; S. Y., J. T., and G. H. wrote the paper.

## Acknowledgments

We thank Prof. Hongning Tong (Institute of Crop Sciences, Chinese Academy of Agricultural Sciences) for kindly providing *osbzr1-1* and *osbzr1-2* seeds. We would like to express our sincere thanks for the strong support of Ningbo Yongxin Optics Scientific Research Instrument (NE950FL) for providing accurate and effective photography service for this research.

