## Supplementary material for "Brassinosteroid-auxin crosstalk shapes rice flag leaf angle to boost dense planting yield": Yu et al SI.pdf

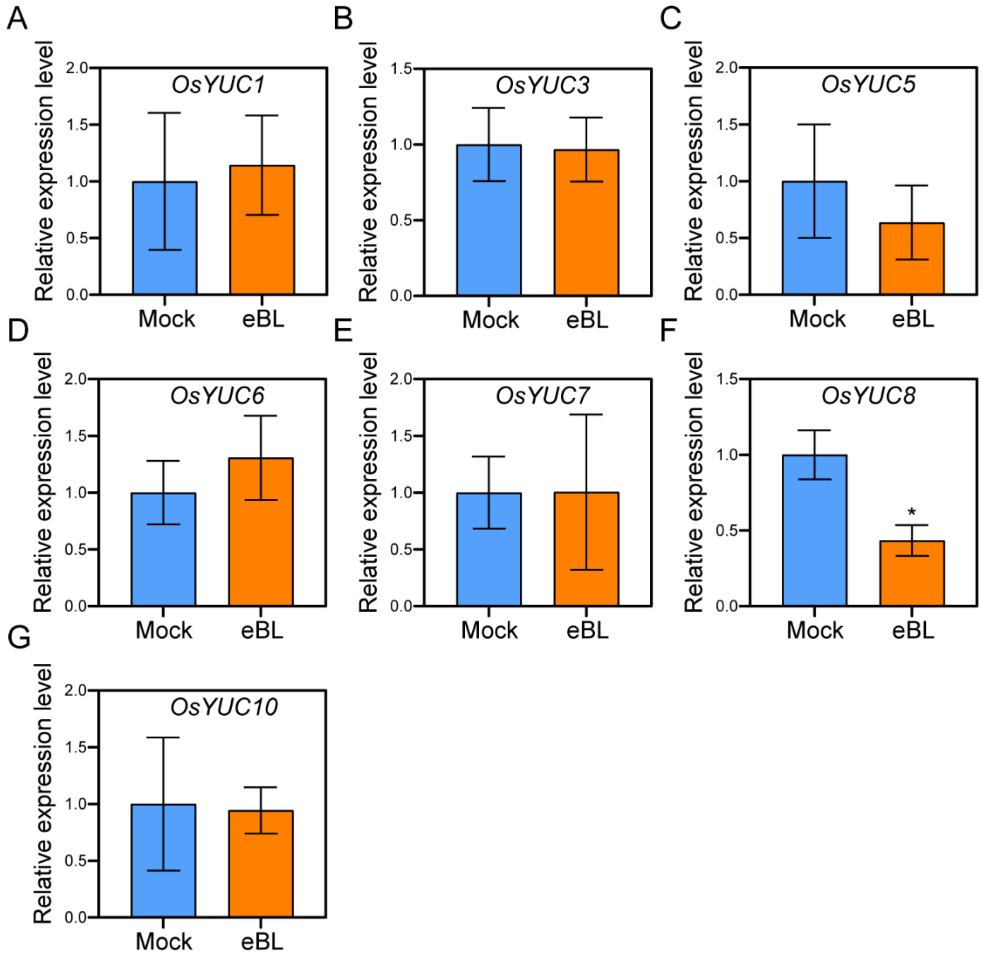


**Supplementary Figure S1. *OsYUC8* expression is inhibited by external eBL treatment.**
**A to G)** Relative expression level of *OsYUC1* **(A)**, *OsYUC3* **(B)**, *OsYUC5* **(C)**, *OsYUC6* **(D)**, *OsYUC7* **(E)**, *OsYUC8* **(F)**, *OsYUC10* **(G)** under eBL treatment. dd$H_2O$ was used as the mock treatment. Error bars are ± SE, n = 3 biological replicates. Student's *t*-test: *$P < 0.05$.

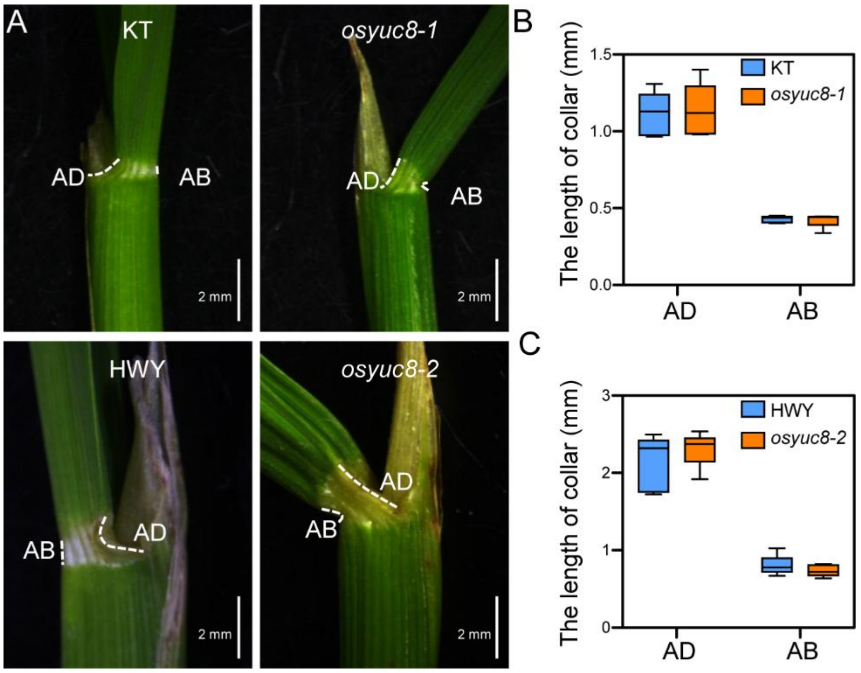


**Supplementary Figure S2. The lamina joint morphology of flag leaf is normal in *osyuc8* mutants.**

**A)** Representative flag leaf lamina joint images in WT and *osyuc8* mutants. The upper two images are KT (WT) and *osyuc8-1*, and the lower two images are HWY (WT) and *osyuc8-2*. AD, adaxial side; AB, abaxial side. The white dashed lines depict the length of AB and AD. Scale bars, 2 mm. **B and C)** Statistical analysis of the lengths of abaxial and adaxial sides shown in **(A)**. n = 5 biological replicates.

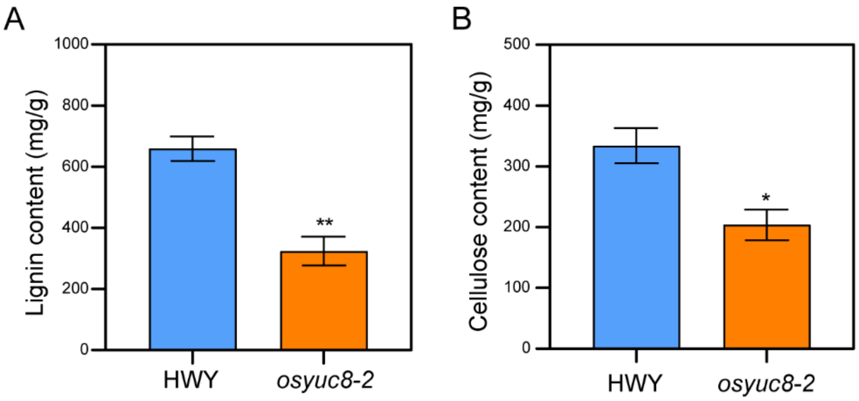


**Supplementary Figure S3. *Osyuc8-2* mutants exhibit decreased lignin and cellulose levels.**
**A)** Statistical analysis of lignin content in the lamina joints of 100-day-old WT and *osyuc8-2* mutants. Error bars are ± SE, n = 4 biological replicates. Student's t-test: $^{**}P < 0.01$.
**B)** Statistical analysis of cellulose content in the lamina joints of 100-day-old WT and *osyuc8-2* mutants. Error bars are ± SE, n = 4 biological replicates. Student's t-test: $^{*}P < 0.05$.

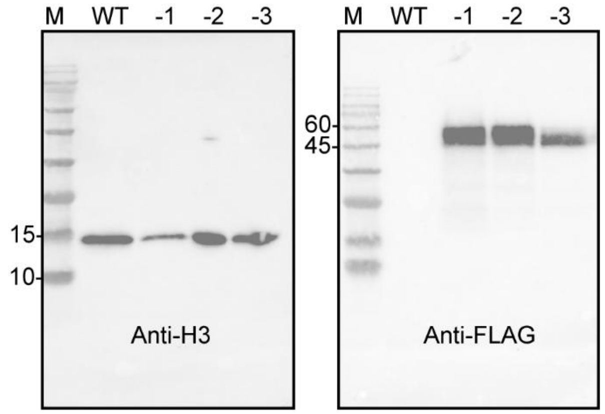


**Supplementary Figure S4. Validation of OsBZR1-FLAG transgenic lines by western blot.** Total proteins were extracted from the flag leaf lamina joints of 100-day-old rice plants. Histone H3 protein was used as an internal reference, and the Flag-tagged OsBZR1 protein was detected using FLAG antibodies. The suffixes -1, -2, and -3 indicate independent transgenic lines exhibiting distinct levels of OsBZR1 accumulation.

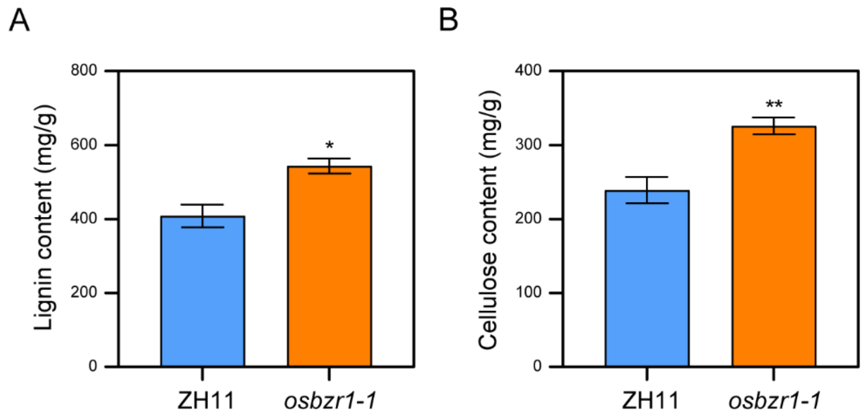


**Supplementary Figure S5.** Increased lignin and cellulose content in *osbzr1-1* mutants.
**A)** Statistical analysis of lignin content in the lamina joints of 100-day-old WT and *osbzr1-1* mutants. Error bars are ± SE, n = 4 biological replicates. Student's t-test: *$P < 0.05$.
**B)** Statistical analysis of cellulose content in the lamina joints of 100-day-old WT and *osbzr1-1* mutants. Error bars are ± SE, n = 4 biological replicates. Student's t-test: **$P < 0.01$.

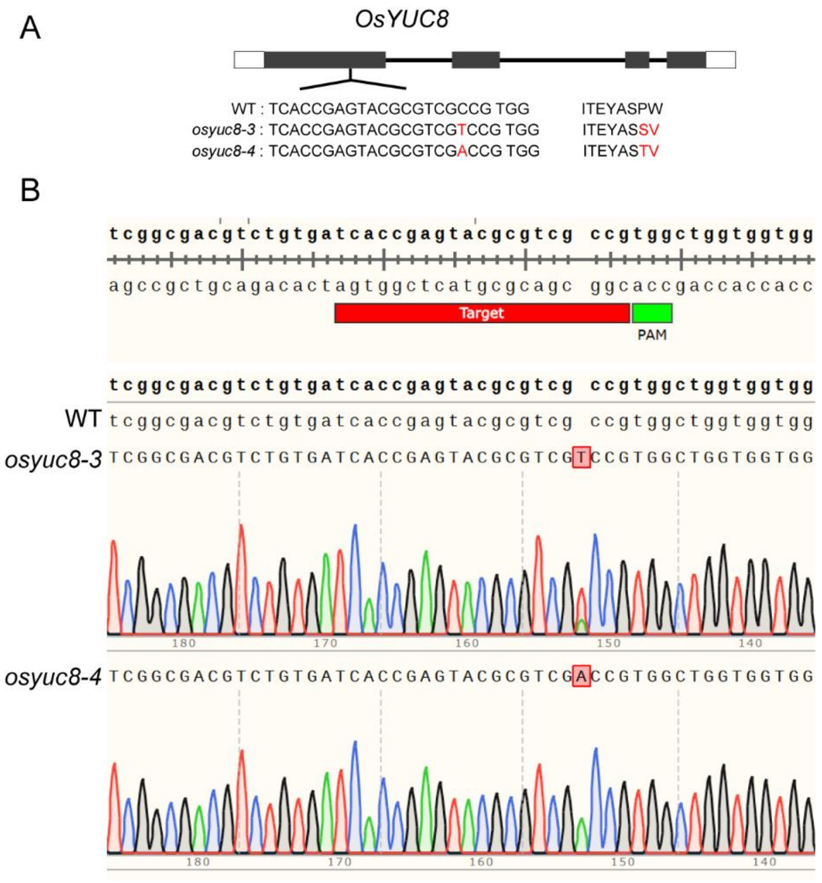


**Supplementary Figure S6.** Knockout *OsYUC8* in *osbzr1-1* background.
**A)** Knockout mutants of *OsYUC8* via CRISPR/Cas9. Black boxes denote exons, and white boxes indicate the 5' and 3' untranslated regions (UTRs). The *osyuc8-3* and *osyuc8-4* mutants carry a single T and A insertion in the first exon, causing a frameshift mutation, respectively.
**B)** Sequencing alignments of *OsYUC8* knockout lines. The *osyuc8-3* and *osyuc8-4* mutants carry a single T and A insertion in the first exon, respectively. Red labels indicate the target sequences recognized by the gRNA. Green labels indicate the protospacer adjacent motif (PAM) sequences required for Cas9 cleavage.

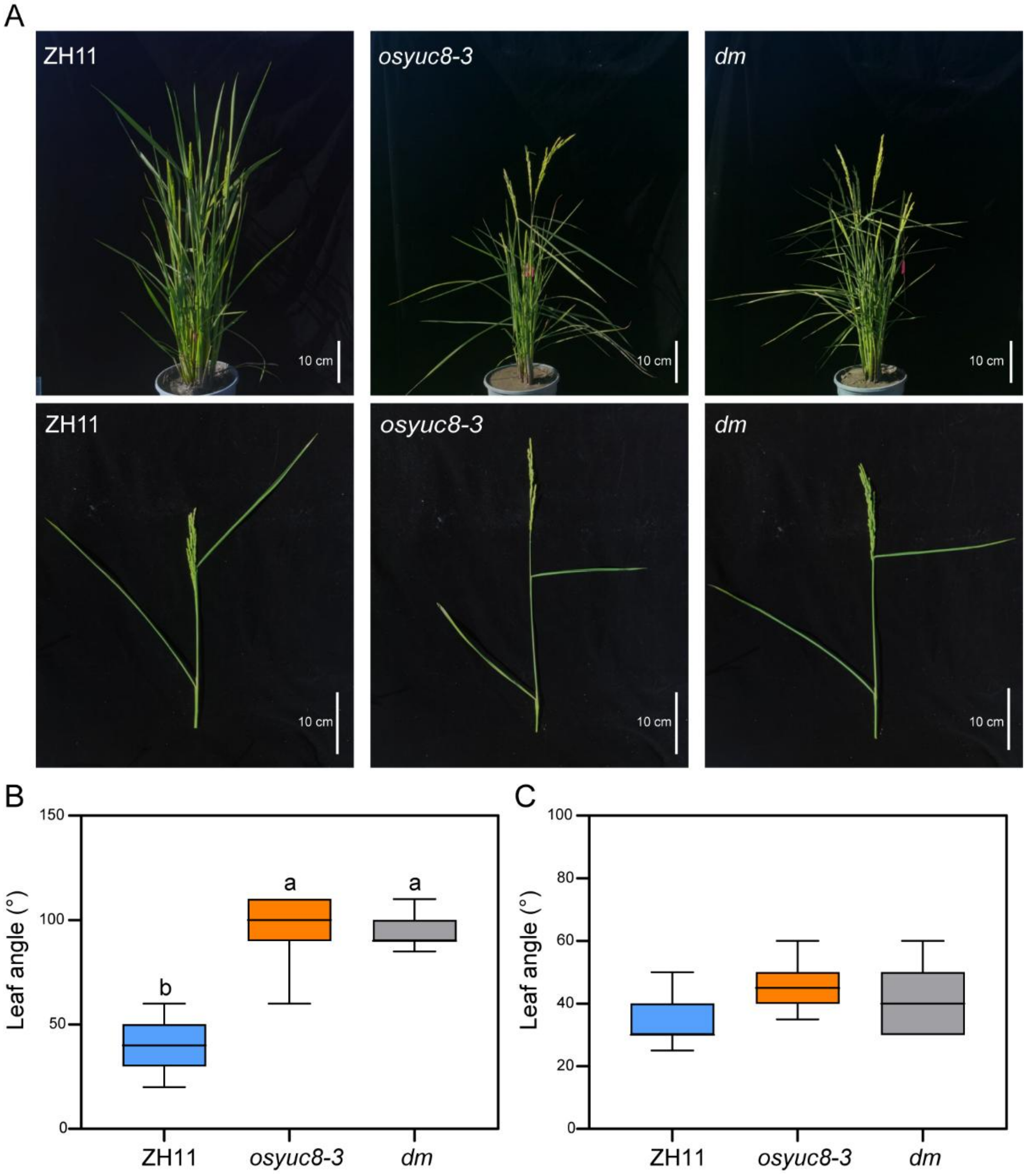


**Supplementary Figure S7.** Genetic epistasis of *OsYUC8* and *OsBZR1* in controlling FLA.
**A)** Representative images of WT, *osyuc8-3*, and *osbzr1-1 osyuc8-3* double mutant (*dm*) grown for 100 days. Scale bars, 10 cm.
**B and C)** Analysis of the FLA **(B)** and the top second leaf angle **(C)** in WT, *osyuc8-3*, and *dm* grown for 100 days. Different letters indicate significant differences, $P < 0.01$ by one-way ANOVA analysis, n = 15 biological replicates.

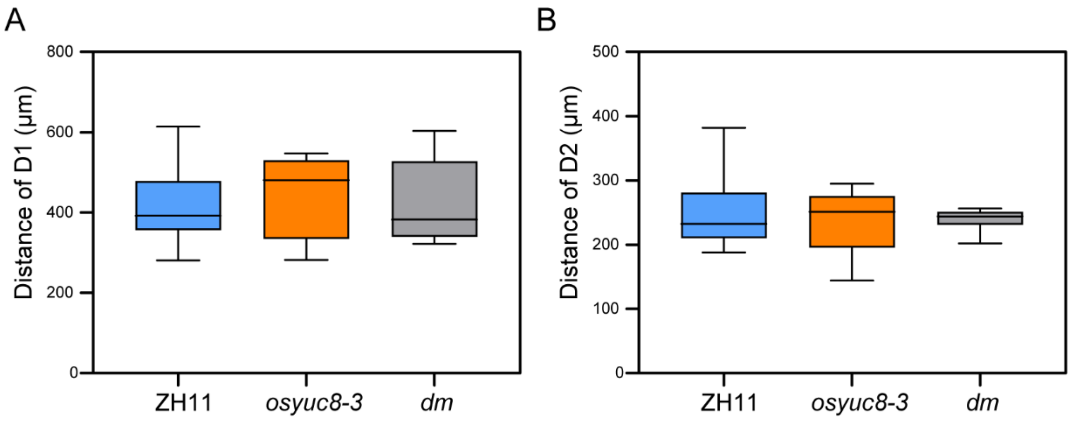


**Supplementary Figure S8. Lamina joint morphology of flag leaf of *osbzr1-1 osyuc8-3* mutant is normal.**

**A)** Quantification of the sclerenchyma cell layer thickness in cross sections of the lamina joint of flag leaf from WT, *osyuc8-3*, and *dm* grown for 100 days. D1, distance of lamina joint abaxial margin to vascular bundles. n = 9 biological replicates.

**B)** Quantification of the parenchyma cell layer thickness in cross sections of the lamina joint of flag leaf from WT, *osyuc8-3*, and *dm* grown for 100 days. D2, distance of lamina joint adaxial margin to vascular bundles. n = 9 biological replicates.

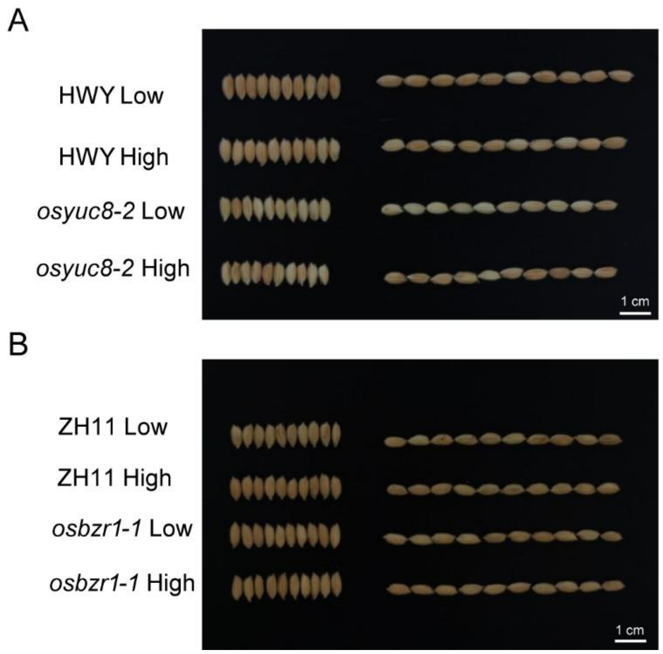


**Supplementary Figure S9. Grain size of WT, *osyuc8-2*, and *osbzr1-1* mutants under different planting densities.**
**A)** Grain length and width of WT (HWY) and *osyuc8-2* mutants under low-density (Low, 25 plants $m^{-2}$) and high-density (High, 49 plants $m^{-2}$) planting. Ten grains were randomly selected from WT and *osyuc8-2* mutants grown under different planting densities and arranged side by side. Scale bars, 1 cm.
**B)** Grain length and width of WT (ZH11) and *osbzr1-1* mutants under low- and high-density planting. Ten grains were randomly selected from WT and *osbzr1-1* mutants grown under different planting densities and arranged side by side. Scale bars, 1 cm.

**Supplementary Table S1.** Primers used in this study

| **Primer Name** | **Primer Sequence** |
|---|---|
| **Primers for genotyping PCR** | |
| *OsYUC8*-F | TGTTTGGGGCCTCGACATTT |
| *OsYUC8*-R | CTGTGGACGACTACACCGAG |
| *OsYUC8*-F2 | TCCGCTACCTCCACTCCTAC |
| *OsYUC8*-R2 | GTCCAGGCACATCTCCATCC |
| *OsBZR1*-F | AGAGGGAAAGCACGCTACTG |
| *OsBZR1*-R | CTCGGGAAGCTCGACGAC |
| **Primers for RT-qPCR** | |
| *Ubiquitin*-F | GAGCCTCTGTTCGTCAAGTA |
| *Ubiquitin*-R | ACTCGATGGTCCATTAAACC |
| *qOsYUC1-F* | AAGAAGGTGTTGGTCGTGGG |
| *qOsYUC1-R* | ATGCCGAACGTGGATAGACC |
| *qOsYUC3-F* | TTTGCTGGGAAGCGTGTTCT |
| *qOsYUC3-R* | CCAAAGGTGGACTGACCCAG |
| *qOsYUC5-F* | AGCTACAAGAGAGGGGACGA |
| *qOsYUC5-R* | GAGGCCAAATGTTGAGATGCC |
| *qOsYUC6-F* | CGGATACCAAAGCAACGTCC |
| *qOsYUC6-R* | GACTCCCCCTTCCAACCATC |
| *qOsYUC7-F* | GCTACCGCAGCAATGTGCC |
| *qOsYUC7-R* | CCCGACTCACCCTTCCATCC |
| *qOsYUC8*-F | CCAACATCTCCTCGGTGTAG |
| *qOsYUC8*-R | GCATCAGACAAGCAACATCC |
| *qOsYUC10*-F | CCACCAAGCAGTGGCTCAAG |
| *qOsYUC10*-R | CTCCAGCGCTTCATCTGCTT |
| *qOsBU1-F* | GAGACGTGCAGCTACATCAAGAG |
| *qOsBU1-R* | TGGGCTGTTGTGATCCATGC |
| *qOsBZR1-F* | TCCCGTACCTGTCATGTGCATC |
| *qOsBZR1-R* | GGTACGTCAAAGCGATCATGCC |
| *qOsIAA20-F* | CGGGATTATTTTGTTCACGTTTC |
| *qOsIAA20-R* | CGAGATTTCATTCGTCATGCTTA |
| **Primers for ChIP-qPCR** | |
| CF1 *OsYUC8* promoter-F | GAGATCGAAAGCGTGTTCCATC |
| CF1 *OsYUC8* promoter-R | CCCCGAGAAACTCGAAGAAC |
| CF2 *OsYUC8* promoter-F | TTGGATCCATTTGCCCTTGC |
| CF2 *OsYUC8* promoter-R | AGAGGCATTTCACCGAACAC |
| CF3 *OsYUC8* promoter-F | ACAAACAGGGCAGCTAAAATTTC |
| CF3 *OsYUC8* promoter-R | TGATGGACACAAAGCAGCTAC |
| CF4 *OsYUC8* promoter-F | AATCCACATAGAGAGGCAGAGC |
| CF4 *OsYUC8* promoter-R | GCAATGGAATGGACACATCTCC |
